# SUMOylation regulates protein cargo in Astrocyte-derived small extracellular vesicles

**DOI:** 10.1101/2020.09.15.298554

**Authors:** Anllely Fernández, Maxs Méndez, Octavia Santis, Katherine Corvalan, Maria-Teresa Gomez, Peter Landgraf, Thilo Kahne, Alejandro Rojas-Fernandez, Ursula Wyneken

## Abstract

Recent studies have described a new mechanism of intercellular communication mediated by various types of extracellular vesicles (EVs). In particular, exosomes are small EVs (sEVs) released to the extracellular environment by the fusion of the endosomal pathway-related multivesicular bodies (containing intraluminal vesicles) with the plasma membrane. sEVs contain a molecular cargo consisting of lipids, proteins, and nucleic acids. However, the loading mechanisms for this complex molecular cargo have not yet been completely elucidated. In that line, the post translational modification SUMO (Small Ubiquitin-like Modifier) has been shown to impact the incorporation of select proteins into sEVs. We therefore decided to investigate whether SUMOylation is a mechanism that defines protein loading to sEVs. In order to investigate the role of SUMOylation in cargo loading into sEVs, we utilized astrocytes, an essential cell type of the central nervous system with homeostatic functions, to study the impact of SUMOylation on the protein cargo of sEVs. Following SUMO overexpression, achieved by transfection of SUMO plasmids or experimental conditions that modulate SUMOylation in primary astrocyte cultures, we detected proteins related to cell division, translation, and transcription by mass-spectrometry. In astrocyte cultures treated with the general SUMOylation inhibitor 2-D08 (2′,3′,4′-trihydroxy-flavone, 2-(2,3,4-Trihydroxyphenyl)-4H-1-Benzopyran-4-one) we observed an increase in the number of sEVs and a decreased amount of protein cargo within them. In turn, in astrocytes treated with the stress hormone corticosterone, we found an increase of SUMO-2 conjugated proteins and sEVs from these cells contained an augmented protein cargo. In this case, the proteins detected with mass-spectrometry were mostly proteins related to protein translation. To test whether astrocyte-derived sEVs obtained in these experimental conditions could modulate protein synthesis in target cells, we incubated primary neurons with astrocyte-derived sEVs. sEVs from corticosterone-treated astrocytes stimulated protein synthesis while no difference was found with sEVs derived from 2-D08-treated astrocytes. Our results show that SUMO conjugation plays a fundamental role in defining the protein cargo of sEVs impacting the physiological function of target cells.

## INTRODUCTION

Extracellular vesicles (EVs), i.e. extracellular structures enclosed by a lipid bilayer, are novel players in intercellular communication (Mathieu, Martin-Jaular, Lavieu, & Théry, 2019; Van Niel, D’Angelo, & Raposo, 2018). EVs that are released after the fusion of multivesicular bodies (MVBs) with the plasma membrane, allowing the secretion of intraluminal vesicles (ILVs) of a diameter from 30 to 160 nm, are termed exosomes. In turn, when EVs are generated directly from the plasma membrane, their diameter varies between 30 nm and 1 μm, and they are called microvesicles. Given the heterogeneity of EV preparations, exosome-containing fractions are now more precisely termed small extracellular vesicles (sEVs) (Colombo, Raposo, & Théry, 2014; Witwer & Théry, 2019). Remarkably, the molecular content (i.e. lipids, proteins and nucleic acids) of sEVs depends on the physiological or pathophysiological state of the donor cell (Mathieu et al., 2019). Regarding proteins, sEVs contain a common protein pattern (or protein markers), such as CD63, CD9, Flotillin or TSG101, as well as a cell type-specific and physiological state-dependent protein cargo (Doyle & Wang, 2019; Jeppesen et al., 2019; Kowal et al., 2016; Zhang, Liu, Liu, & Tang, 2019).

The transfer of sEVs to target cells can regulate the recipient cell function in a cargo-dependent manner, e.g. influencing the development or progression pathophysiological processes (Isola & Chen, 2016; Kalluri & LeBleu, 2020). It is therefore of great relevance to understand precisely the mechanisms involved in the loading of biomolecules, which will define the biological effect of sEVs.

The mechanisms that participate in the biogenesis of sEVs may act independently or in coordination with the classification/loading of molecules into the vesicles (Simons & Raposo, 2009). The first mechanism described depends on the endosomal sorting complex required for transport (ESCRT), composed of 4 multiproteic complexes (0 to III). Secondly, ceramides constitute a primary lipid factor with physical and structural properties that facilitate the biogenesis of intra-luminal vesicles (Holopainen, Angelova, & Kinnunen, 2000; Simons & Raposo, 2009). The third mechanism is through tetraspanins, which are capable of modulating the formation of membrane microdomains, playing a fundamental role not only in the formation of vesicles, but also in the selection of proteins that are specifically incorporated into them (Zhang et al., 2019).

Post-translational modifications that contribute to the selective loading of proteins into sEVs include protein modification by ubiquitin and ubiquitin-like proteins (UBLs) such as SUMO (Small Ubiquitin-like Modifier) (Ageta & Tsuchida, 2019; Colombo et al., 2014). SUMO conjugation to lysine residues impacts on the function of proteins and importantly, the formation of multi-protein complexes (J. M. Desterro, Thomson, & Hay, 1997; Johnson & Blobel, 1997). Moreover, SUMO can interact in a non-covalent manner to SUMO interactive motif, also known as SIM (Hecker, Rabiller, Haglund, Bayer, & Dikic, 2006; Song, Durrin, Wilkinson, Krontiris, & Chen, 2004). There are three functional SUMO homologs in mammals with a molecular weight of ∼10KDa: SUMO-1, SUMO-2, SUMO-3 (J. M. P. Desterro, Rodriguez, Kemp, & Ronald T, 1999; Liang et al., 2016; Melchior, 2000). SUMOylation regulates the activity, stability, and sub-cellular localization of proteins, primarily modifying protein-protein interactions (Geiss-Friedlander & Melchior, 2007). SUMO modification is involved in a wide variety of cellular processes, affecting among others translation, transcription, replication, chromosome segregation, DNA repair, differentiation, apoptosis, senescence, cell cycle, nuclear transport and signal transduction (Hay, 2005; Hendriks & Vertegaal, 2016; Khan, Pandupuspitasari, Huang, Hao, & Zhang, 2016; Pichler, Fatouros, Lee, & Eisenhardt, 2017).

This modification affects the ability of the ribonucleoprotein hnRNPA2B1 to export microRNAs (miRNAs) in sEVs (Villarroya-Beltri et al., 2013). Moreover, α-synuclein is SUMOylated in the sEV lumen while GFP is sorted into sEVs only when it is conjugated with SUMO-2 (Kunadt et al., 2015). In the same context, in homogenized primary astrocyte cultures, the glycolytic enzyme Aldolase C (ALDOC) is detected with two molecular weights: 36KDa (expected weight) and 55kDa (the putatively SUMO-conjugated form). Importantly, in rat serum sEVs, only the high molecular weight form of the enzyme can be found suggesting that SUMOylation determines the loading of this enzyme (Gómez-Molina et al., 2019). In summary, these results strongly support the concept that SUMO conjugation of proteins might dynamically regulate the classification and loading of proteins in sEVs. Previously, the effect of SUMOylation as a general loading mechanism in sEVs had not been described, neither were the proteins SUMOylated within sEVs. In the present work, we describe that SUMOylation regulates the protein cargo of sEVs derived from HeLa cells and astrocytes. Identification of proteins by mass spectrometry indicates that after stimulation of SUMOylation in donor cells (by SUMO overexpression or corticosterone treatment), proteins related to cell division, transcription and translation are enhanced in sEVs. Moreover, sEVs derived from corticosterone-treated donor astrocytes increase *de novo* protein synthesis in target neurons.

## METHODOLOGY

### Reagents and antibodies

αALDOA (#390733), αALIX (#53540), αEF-2 (#166415), αSUMO-1 (#5308) were purchased from Santa Cruz (Dallas, Texas). αFLOTILLIN (#610820) and αGM130 (#610823), were purchased from BD Transduction Labs (New Jersey, USA). Goat anti mouse (#926-80010), Goat anti rabbit (#926-80011) were purchased from Licor (Nebraska, USA). Donkey anti Sheep (#16041), Alexa Fluor 488 mouse (#A21202), DMEM medium (#12100046), Fetal bovine serum (#26140079), G418 (#11811023) Penicillin/Streptomycin (#15140122), neurobasal medium (#21103049), B27 (#17504044), OptiMEM (#31985062), Lipofectamine (#11668030), PBS (#14190), Total Exosome Isolation kit (#4478359), chemiluminescence kit (#32106), DMSO (#85190), IL-1β (#14-8018-62), MOWIOL (#81381) were purchased from Thermo Fisher (Massachusetts, USA). N-Ethylmaleimide (NEM, #34115), DAPI (4′,6-diamidino-2-phenylindole; D9542), Poly-D-Lysine (#4174), Corticosterone (#27840), 2-D08 (#1052) were purchased from Sigma (Missouri, USA). MAP2 (#MAB378) was purchased from Millipore (Massachusetts, USA). αGFAP (#G2032) were purchased from US Biological (Massachusetts, United States). Cheap αSUMO-2 was generously donated by Dr. Ronald T. Hay, University of Dundee, UK.

### Cell culture

HeLa cells were maintained in DMEM containing 10% Fetal bovine serum with 100 units/ml of penicillin and 100 μg/ml of streptomycin and 200 μg/ml of G418 (for maintaining stable cell lines) incubated at 37°C, with 5% CO2 and 95% humidity. Astrocytes were obtained from the telencephalon of post-natal rats (at postnatal day 1) as already described (Luarte et al., 2020). Astrocytes were maintained in DMEM containing 10% Fetal bovine serum with 100 units/ml of penicillin and 100 μg/ml of streptomycin incubated at 37°C, with 5% CO2 and 95% humidity. On days 4 and 8 *in vitro* (DIV), a total change of medium was made. After 15 DIV, the astrocytes were re-plated to decrease the presence of microglia and seeded at a confluence of 70-80%. Cortical neurons of the rat embryo brain (E18) were dissociated from the cerebral cortex and 20,000-40,000 cells were seeded on coverslips coated with Poly-D-Lysine using 35mm plates. Cortical neurons were maintained *in vitro* for 15 days in neurobasal medium supplemented with B27 and 100 units/mL of penicillin and 100 μg/mL of streptomycin incubated at 37°C, with 5% CO2 and 95% humidity. Cells were cultured in plates until they reached 60% confluence or after 15DIV (astrocytes and cortical neurons). At that time, some wells were transfected or treated with different reagent, as indicated in each case. At the end of these treatments, cells were trypsinized, resuspended in phosphate-buffered saline (PBS) and an aliquot was counted using a Neubauer chamber.

### Transfections

Transfections were performed after reaching 60% cell confluence using a 3:1 ratio of DNA: Lipofectamine in Opti-MEM. pCDNA3.1 HIS-SUMO-1 and HIS-SUMO-2 plasmid were generously donated by Dr. Ronald T. Hay, University of Dundee, UK.

### Obtaining sEVs

Cell cultures were grown in a sEV free culture medium for 72 hours (DMEM with 10% FBS, depleted by ultracentrifugation for 2 hours at 100,000xg). Then, the conditioned media of these cells was harvested to isolate sEVs by ultracentrifugation. sEVs were isolated by serial centrifugations as already described (Luarte et al., 2020; Théry, Amigorena, Raposo, & Clayton, 2006): After 30 minutes at 2,000g the supernatant was recovered, the centrifuged for 45 minutes at 10,000g. Then the supernatant was centrifuged for 2 hours at 100,000g, the supernatant was eliminated and the sEV enriched pellet was washed and re-suspended in PBS and stored at −80°C until use. Or, using Total Exosome Isolation kit (Thermo-Fisher), the sEV containing fraction was obtained by centrifugation for 30 minutes at 2,000g. The supernatant was resuspended with 0.5 volumes of the Total Exosome Isolation solution and incubated overnight at 4°C. Then the mix was centrifuged for 1 hour at 10,000g and the pellet was resuspended in PBS and stored at −80°C until use.

### Nanoparticle tracking analysis

The nano particle tracking analysis (Nanosight NS300, Malvern Instruments, Malvern, UK) was used to determine particle concentration and size distribution. Samples were diluted 5 or 10 times in PBS to obtain ≥80 particles per field for analysis and 3 videos were recorded with a duration of 30 seconds each.

### Western blots

Cells were lysed in RIPA buffer (150mM NaCl, 25mM Tris-HCL pH 7.4, NP-40, 0.5% sodium deoxycholate, 20mM NEM (N-Ethylmaleimide)). Protein quantification of the homogenates and sEVs (sEV resuspended with with 0.1% SDS) were determined using the BCA method (Smith et al., 1985). The samples were boiled with loading buffer and 20mM NEM (N-ethyl-maleimide, modify cysteine residues in proteins and peptides) for 5 minutes at 100°C. Each lane was loaded with either the same amount of total protein or the same number of vesicles, as depicted. Proteins were separated in 12% polyacrylamide gels under denaturing conditions. Gels were stained with Coomassie dye or transferred to nitrocellulose membranes. The membranes were incubated with blocking solution (5% nonfat milk in PBS) for one hour, then incubated over night with primary antibody. The membranes were washed with PBS. Finally, they were incubated with secondary antibody for 45 minutes and then visualized with a chemiluminescence kit (Pierce ECL #32109).

### SIM-SUMO pulldown

SIM-HALO resins consisting of the SIM sequence fused with a HALO tag and linked to a resin were incubated with 200 μg of sEV protein with RIPA buffer and 20mM NEM for 16 hours at 4°C. They were centrifuged at 500xg for 5 min and washed 8 times with RIPA with 20mM NEM. The supernatant was removed each time. Finally, the pellet was resuspended with loading buffer and 20mM NEM, the samples were boiled, and centrifuged at 10,000xg for 10 minutes and then analyzed by SDS-PAGE followed by Coomassie staining. 10% of the Input eluates were loaded.

### Proteomics

sEV proteins were separated using polyacrylamide gradient gel electrophoresis. Each lane was divided into 8 sections to perform in-gel digestion. Liquid chromatography followed by tandem-mass spectrometry (MS/MS) of the sample fractions was performed on a hybrid dual-pressure linear ion trap/orbitrap mass spectrometer (LTQ Orbitrap Velos Pro, Thermo Scientific) equipped with an EASY-nLC Ultra HPLC (Thermo Scientific). Peptide samples were dissolved in 10 μL of 2% acetonitrile/0.1% trifluoric acid and fractionated on a 75-μm i.d., 25-cm PepMap C18-column, packed with 2 μm of resin (Dionex, Germany). Separation was achieved by applying a gradient of 2% to 35% acetonitrile in 0.1% formic acid over 150 minutes at a flow rate of 300 nL/min. The LTQ Orbitrap Velos Pro MS was exclusively used for CIDfragmentation when acquiring MS/MS spectra, which consisted of an orbitrap full mass spectrometry (MS) scan followed by up to 15 LTQ MS/MS experiments (TOP15) on the most abundant ions detected in the full MS scan. The essential MS settings were as follows: full MS (resolution, 60,000; mass to charge ratio range, 400–2000); MS/MS (Linear Trap; minimum signal threshold, 500; isolation width, 2 Da; dynamic exclusion time setting, 30 seconds; and singly charged ions were excluded from the selection). Normalized collision energy was set to 35%, and activation time was set to 10 milliseconds. Raw data processing and protein identification were performed by ProteomeDiscoverer 1.4 (Thermo Scientific) and a combined database search used the Sequest and Mascot algorithms. The false discovery rate was calculated by the Percolator 2.04 algorithm and was set to <1%. An Ingenuity Pathway Analysis was used for network analysis of functional interactions of proteins (Gómez-Molina et al., 2019). The bio-informatic analysis was done by DAVID Bioinformatics Resources 6.8. pValue was represented as −Log_10_.

### Incubation of target cells with sEVs

A total of 1000 vesicles was added per cell, and this was repeated 24 hours later. Then, 48 hours later, the cells were used for the respective assays: MTT (3-(4,5-Dimethylthiazol-2-yl)-2,5-Diphenyltetrazolium Bromide), DAPI (4′,6-diamidino-2-phenylindole), PI (Propidium iodide) or FUNCAT assay (fluorescence non-canonical amino acid tagging).

### Donor cell treatments

The culture medium was completely changed to sEV free culture medium and the following treatments added 1 hour later: 10ng/ml interleukin 1β (IL-1β), 1μM Corticosterone (CORT) or 30μM 2-D08 (inhibitor of the enzyme UBC9, SUMO-conjugating enzyme) or DMSO (as a control). This was repeated 24 and 48 hours later, and the conditioned medium was collected 72 hours later. The medium was collected to isolate sEVs and the cells were homogenized and stored at −80°C until further use.

### Morphological analysis

Neurons incubated with astrocyte-derived sEVs obtained in the different experimental conditions were analyzed using Sholl analysis using plugins from ImageJ software (National Institute of Health, USA). The dendritic tree was examined in 3 μm increments. The following parameters were obtained: total dendrite length (i.e., the largest radius at which there is an intersection with a neuronal process) and total number of intersections (i.e., the sum of all intersections with each different radius) (Luarte et al., 2020).

### FUNCAT (fluorescent click chemistry) and immunefluorence

To visualize newly synthesized proteins in cells, the FUNCAT method was used according to Daniela Dieterich (Dieck et al., 2012). Cultures were incubated with the non-canonical amino-acid AHA (L-azidohomoalaine), which was incorporated into newly synthesized proteins instead of methionine. As a negative control, methionine was used. Then, the neurons were fixed and the FUNCAT assay was performed. Briefly, cells were washed with cold PBS-MC (1 mM MgCl_2_, 0.1 mM CaCl_2_ in PBS pH7.4) and directly fixed with 4% paraformaldehyde for 30 minutes, then washed 3 times with PBS pH 7.4 and incubated with B-Block solution (10% normal horse serum, 5% sucrose, 2% BSA, 0.2% TritonX-100 in PBS pH 7.4) for 1 hour at room temperature. Cells were newly washed 3 times with PBS pH 7.8 and incubated overnight with FUNCAT solution (0.2mM Triazole ligand (Tris[(1-benzyl-1H-1,2,3-triazol-4-yl)methyl]amine), 0.5mM TCEP [Tris-(2-carboxyethyl)phosphine hydrochloride], TAMRA (red-fluorescent tetramethylrhodamine) Alkyne tag, 40 μg/ml CuSO_4_ in PBS pH 7.8). The cells were then washed with PBS pH 7.8 and then with PBS pH 7.4 and incubated with primary antibody in B-block+ 0.2% TritonX-100 for 2 hours at room temperature, then were washed with PBS pH 7.4 and incubated with secondary antibody in B-Block solution for 1 hour at room temperature. Finally, the cells were incubated for 10 min with 300mM DAPI in PBS 7.4, washed with PBS pH 7.4 and mounted using MOWIOL mounting media. The images were taken with a confocal LSM 800 Zeiss microscope. The FUNCAT signal was visualized as a fire lookup table, to visually favor the range of expression of the newly synthesized proteins through a range of colors from blue to white, where blue is the absence of new protein synthesis.

### Statistical analysis

One-way ANOVA were performed followed by a post-hoc Tukey test using GraphPad Prism version 5 for Windows, GraphPad Software, La Jolla California USA.

## RESULTS

### SUMOylation in astrocytes leads to a differential cargo in the derived sEVS

Our initial studies were to characterize the vesicles obtained by two methods of sEV purification, the ultracentrifugation method (UC) and a commercial vesicle isolation kit (IK). Both methods produce samples with a similar protein pattern observed in the Coomassie staining. Western blots revealed that the sEV marker protein Alix was enriched in the vesicles compared to the donor astrocytes using both methods. Flotillin in turn was present, but not enriched, while a protein that is not contained in sEVs, GM130, was not detected in the sEV fraction (Figure 1A). Using either method nanoparticle tracking analysis (NTA) showed a similar particle size profile obtaining a mean of 152 nm in the case of UC (Figure 1B) and of 156 nm after using the IK (Figure 1C) in the sEV fractions isolated.

**Figure 1.**
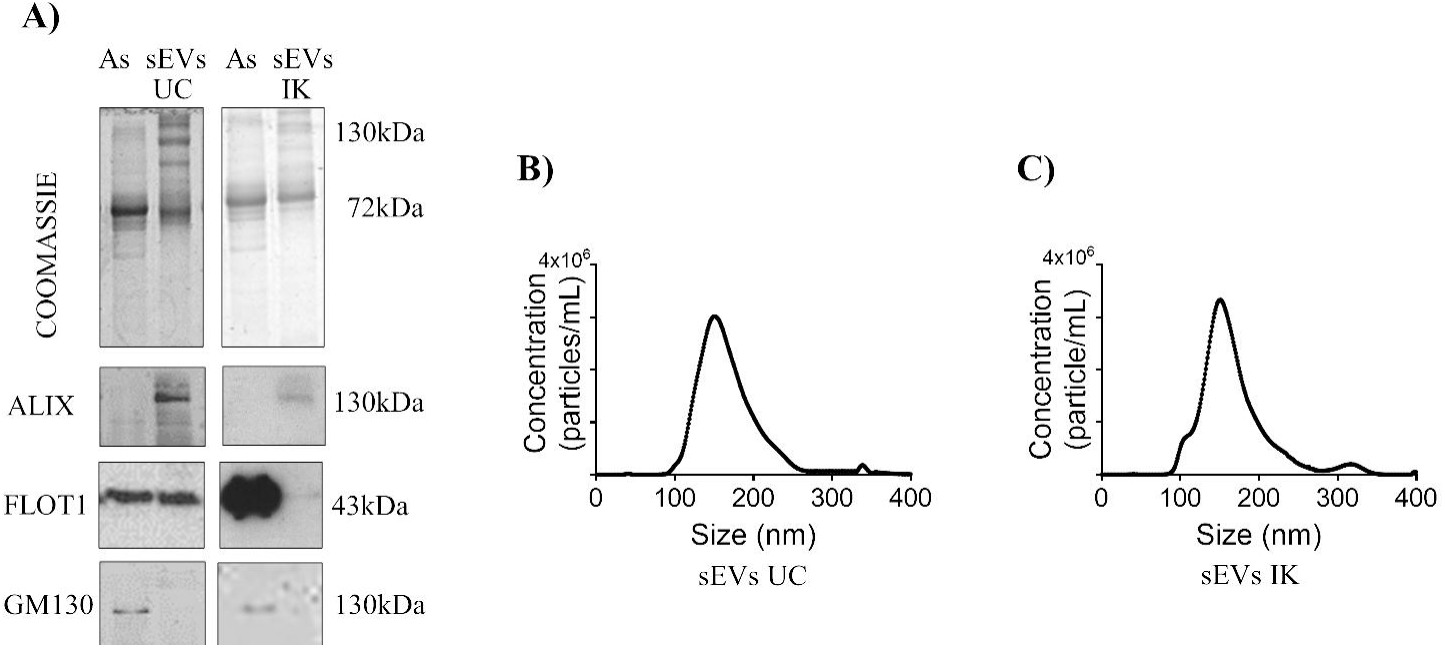
Characterization of small Extracellular Vesicles (sEVs) from astrocytes. A) Coomassie staining and sEVs-enriched proteins detected by Western blot of sEVs isolated by Ultracentrifugation (UC) or sEV isolation kit (IK) compared to donor astrocytes (As). B) sEV size distribution using Nanoparticle Tracking Analysis of sEVs isolated by UC, 152 ± 7.8 nm (mean ± SEM). C) sEV size distribution using sEV IK, mode 156 ± 10.7 nm (mean ± SEM). n= 6 sEVs and cell culture from different murine preparation, 3 technical repeats for each n.

To evaluate the consequences of SUMOylation on the protein content of astrocyte-derived sEVs, transfections of astrocytes with HIS-SUMO-1 (S1) or HIS-SUMO-2 (S2) were carried out, using GFP as a transfection control. We obtained a transfection efficiency of 10-15% by lipofection, evaluated by the detection of GFP-positive cells (data not shown). SUMO conjugation in astrocytes was evaluated by Western blot (Supplementary figure 1A), and in both cases a slightly increased SUMOylation was observed, but do not significant (Supplementary figure 1B and 1C). The UC derived sEVs were subjected to Coomassie staining (Figure 2A) and analyzed by NTA (Figures 2B and 2C). No differences were found in the mean size and sEV yield. At that time, the protein content (in ng) per vesicle was calculated (Figure 2D), and an increase in the protein load in S2 astrocyte-derived sEVs was found. Then, sEV proteins were identified by mass spectrometry (Figure 3 and Supplementary table 1). The Venn and subsequent GO analysis using the DAVID database indicate that the 512 common proteins were significantly enriched in classical exosome components (Figure 3A), confirming that the analyzed fractions are enriched in sEVs (data not shown). When exclusive proteins among conditions were compared, an enrichment in biological processes such as transcription and cell division were found in S1 and S2 sEVs (Figure 3B). To directly identify SUMOylated proteins, we performed a pulldown (PD) of astrocyte-derived sEVs with SIM-HALO resins. Proteins contained in the sEVs (input) and proteins bound to SIM resin were shown by Coomassie staining (Figure 3C). A prominent band was observed over 26kDa in pulled down samples (SIM-bound). In the samples that did not bind to SIM-beads, a banding pattern similar to the input was found at high molecular weights. The samples obtained in the PD were analyzed by mass spectrometry (Supplementary table 1) and we identified an enrichment in biological processes such as Chromatin organization, mRNA processing and cell division (Figure 3D).

**Figure 2.**
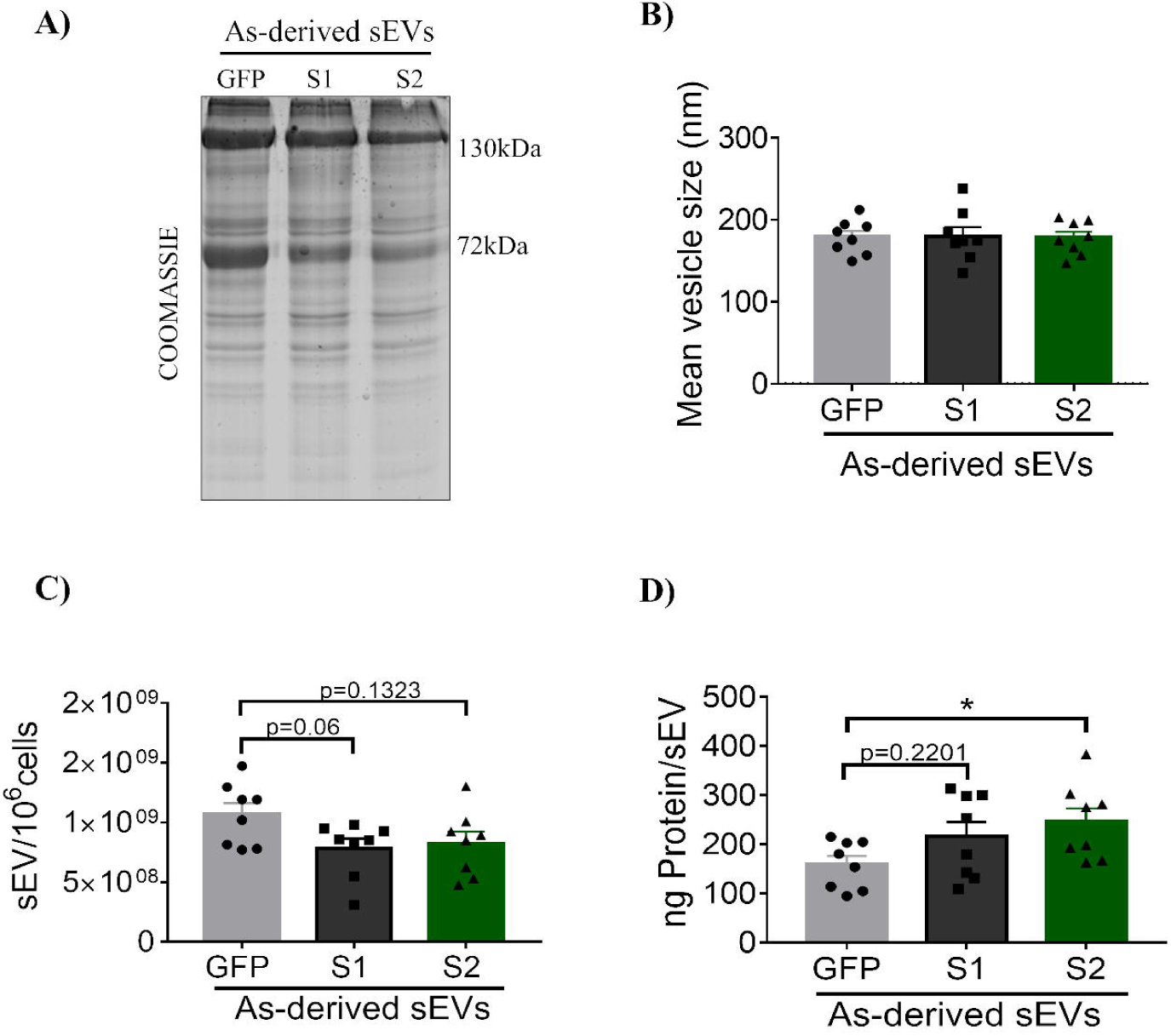
Increased protein loading in Small Extracellular Vesicle (sEVs) of SUMO-2 astrocytes. A) Coomassie staining of the corresponding sEVs. B) Mean sEV size in nm C) Quantification of the number of vesicles obtained from the conditioned media of one million cells. D) Quantification of protein content per sEV. All data are expressed as mean ± SEM and for quantification one-way ANOVA was performed, followed by a post-hoc Tukey test. Statistical significance is represented with asterisks. n = 7-8, sEVs and cell culture from different murine preparation, 3 technical repeats for each n. *p<0.05.

**Figure 3.**
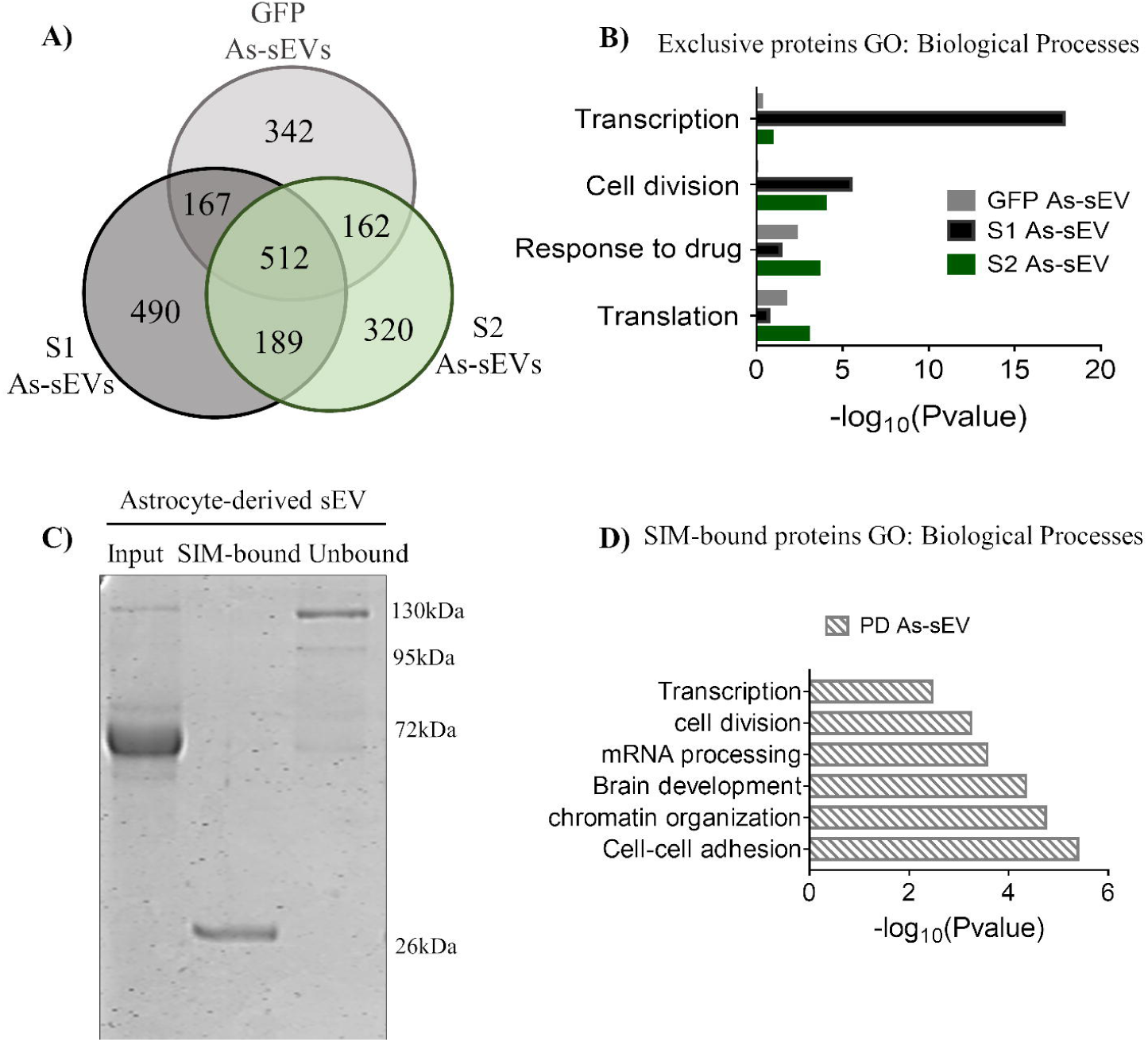
Proteomics of small Extracellular Vesicles (sEVs) from astrocytes expressing SUMO reveal proteins related to transcription, translation and cell division. A) Venn analysis of sEV proteins from astrocytes transfected with GFP, SUMO-1 (S1) or SUMO-2 (S2), identified by mass spectrometry. B) Comparison between biological processes present in exclusive proteins in each sample analyzed by the DAVID database. C) Coomassie staining of 10% input of astrocyte-derived sEVs bound to SIM-beads. D) Biological processes present in proteins bound to SIM-beads analyzed by the DAVID database. Data obtained from n=2-3 6 sEVs from different murine preparation.

Next, a metabolism-related protein ALDOLASE A, (ALDOA) and protein synthesis implicated protein Eukaryotic elongation factor 2 (EEF-2) were validated by Western blot. Equal amounts of protein were loaded per lane in the case of cell samples, while an equal number of vesicles was used in the case of sEV samples. (Supplementary figure 2A). ALDOA in cell samples was found with a molecular weight of ∼36kDa, corresponding to the predicted molecular weight interestingly. In sEVs, ALDOA was observed at ∼55kDa, corresponding to possible SUMOylated form(s), in S1 and S2-derived sEVs (Supplementary figure 2C). Another band is observed at a weight greater than 250 kDa, which could be compatible with a polySUMOylated form of ALDOA. This form is consistently elevated in sEVs from S1 astrocytes (Supplementary figure 2B). In turn, EEF-2 was detected at the predicted molecular weight. This protein tends to have a higher content in sEVs from S1 astrocytes (Supplementary figure 2D).

### Modulation of SUMOylation in astrocytes and-derived sEVs under pathophysiologic-like conditions

In order to change the environmental conditions of sEV donor cells, astrocytes were incubated either with IL-1β, a powerful pro-inflammatory cytokine reported to stimulate SUMOylation (Hajmrle et al., 2014; Miranda, Loeser, & Yammani, 2010), or with corticosterone as an *in vitro* stress model, because stress *in vivo* enhances levels of SUMOylated proteins in sEVs (Gómez-Molina et al., 2019). In the opposing direction, SUMOylation was inhibited with 2-D08, an inhibitor of the enzyme E2 (UbC9) (Kim, Keyser, & Schneekloth, 2014).

Astrocytes were treated one time per day for 3 days with 10ng/ml IL-1β, 1μM Corticosterone (CORT), 30μM 2-D08 or DMSO as control. We first analyzed the SUMOylation pattern of proteins by Western blot (Figure 4A). Quantification showed that corticosterone increased the conjugation of proteins with SUMO-2, while 2-D08 decreased protein conjugation with SUMO-1 and SUMO-2 (Figure 4B and C). To check whether astrocytes were affected after the treatments, the astrocyte marker protein glial fibrillary acid protein (GFAP) was detected in these cultures: 2-D08 decreased GFAP immunoreactivity, and a recovery of its levels was obtained after corticosterone plus 2-D08 (Figure 4D and E).

**Figure 4.**
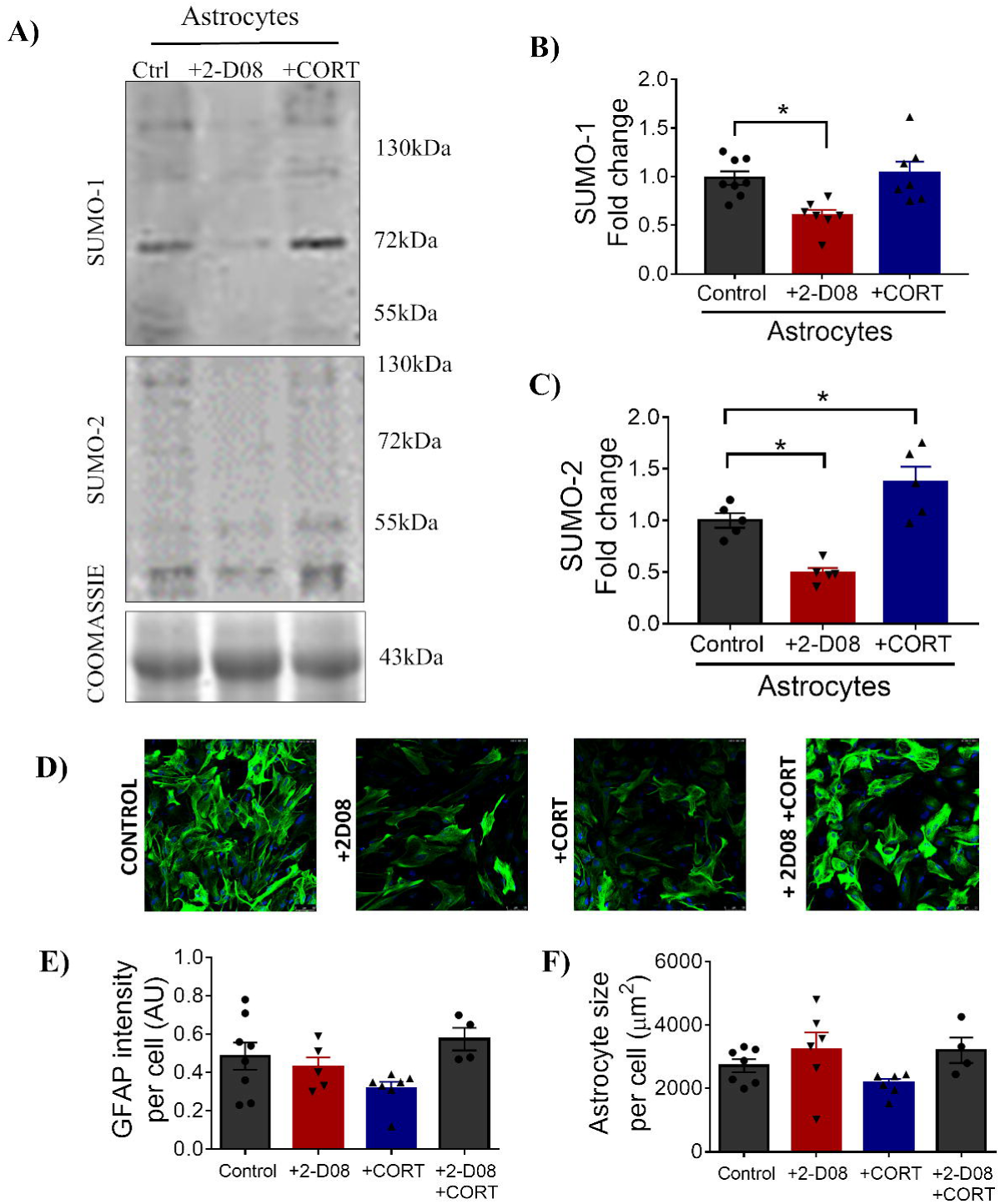
Treatment with 2-D08 decreases and corticosterone increases SUMOylation in astrocytes. Astrocytes were treated 3 times with DMSO (control), 10 ng/ml IL-1β, 1μM corticosterone (CORT) or 30 μM 2-D08 (UBC9 inhibitor), in 24-hour intervals. sEVs were prepared 24 hours after the last treatment. A) Detection by Western blot of proteins conjugated with SUMO-1 or SUMO-2 in treated cells. B) Quantification of protein conjugated with SUMO-1 by densitometry, expressed as fold change with respect to the control and corrected by the densitometry of Coomassie staining. C) Quantification of protein conjugated with SUMO-2 in the same conditions. D) Astrocytes were stained with GFAP antibody (green) and DAPI (blue). E) Quantification of the intensity of GFAP signal. F) Quantification of astrocyte size. All data are expressed as mean ± SEM and for quantification one-way ANOVA was performed, followed by a post-hoc Tukey test. Statistical significance is represented with asterisks. n = 5-8, sEVs and cell culture from different murine preparation, 3 technical repeats for each n. *p<0.05).

Then, we studied the possible effects of these sEVs, isolated by IK, on target cells. First, the same number of vesicles were loaded in each lane of an SDS-PAGE gel to observe the general protein pattern (Figure 5A). As confirmed later, the 2 D08-derived sEVs contained less proteins. There were no differences in the average size of the vesicles derived from the three experimental conditions, as revealed by NTA (Figure 5B). Subsequently, the number of vesicles released per one million cells was quantified. The total number of cells was quantified at the time of sEV collection (Figure 5C). A strong increase of sEVs released from cells treated with 2-D08 was observed, while a decrease in the amount of proteins contained per sEV was observed after 2-D08 treatment (Figure 5D). This result is compatible with the previous Coomassie staining. Interestingly, in HeLa cells a similar effect of 2-D08 was found (Supplementary figure 3).

**Figure 5.**
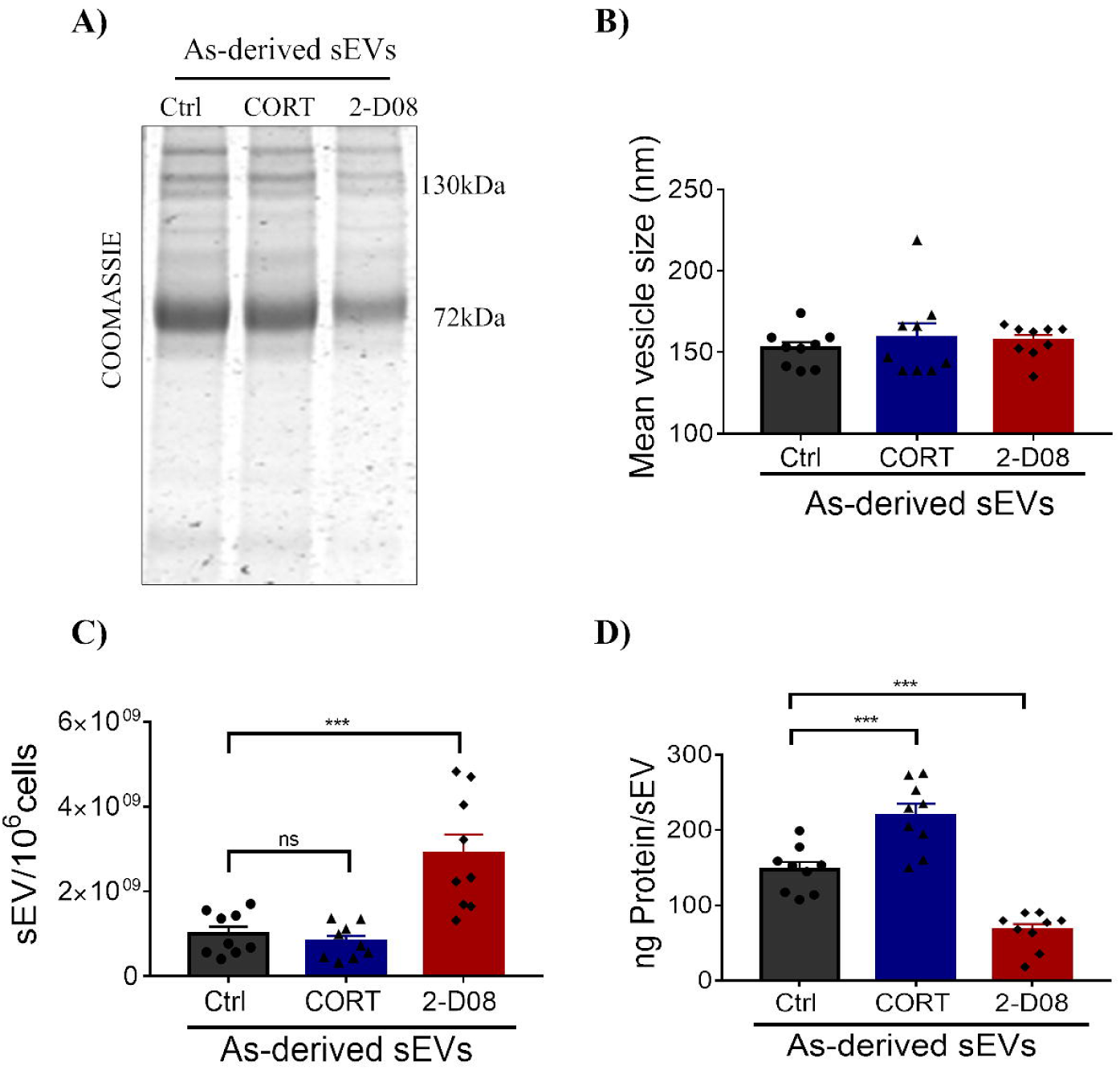
SUMOylation inhibition prevents, and corticosterone increases protein load in small Extracellular Vesicles (sEVs) from Astrocytes. Astrocytes were treated 3 times with DMSO (control), 10 ng/ml IL-1β, 1μM corticosterone (CORT) or 30 μM 2-D08 (UBC9 inhibitor), in 24-hour intervals. sEVs were prepared 24 hours after the last treatment. A) Coomassie staining shows the sEV proteins contained in 10 million sEVs. B) Mean sEV size in nm. C) Number of vesicles released per million cells. D) Quantification of protein content (in ng) per sEV. All data are expressed as mean ± SEM and for quantification one-way ANOVA was performed, followed by a post-hoc Tukey test. Statistical significance is represented with asterisks (n = 9, sEVs and cell culture from different murine preparation, 3 technical repeats for each n.***p<0.001).

### Effect of astrocyte-derived sEVs on protein synthesis in neurons

We next identified the sEV protein content by mass spectrometry derived from control, corticosterone or 2-D08 treated astrocytes (Figure 6A and Supplementary table 3). In total, 149 exclusive proteins were identified in sEVs from corticosterone treated astrocytes. Of these, 19 were related with protein synthesis (Table 1). Comparing protein synthesis-related hits (biological processes) among conditions, an enrichment in rRNA transcription and transcription by RNA polymerase I was found in sEVs from corticosterone-treated astrocytes. Then, we could predict possible SUMOylation sites in 15 of these proteins using the GPS-SUMO database (Zhao et al., 2014).

**Table 1.** **List of the proteins involved in protein synthesis in sEVs derived from astrocytes treated with corticosterone (CORT).** K (Lysine) is a possible site of SUMOylation. Data obtained from n=4-6 biological repeats.

**Figure 6.**
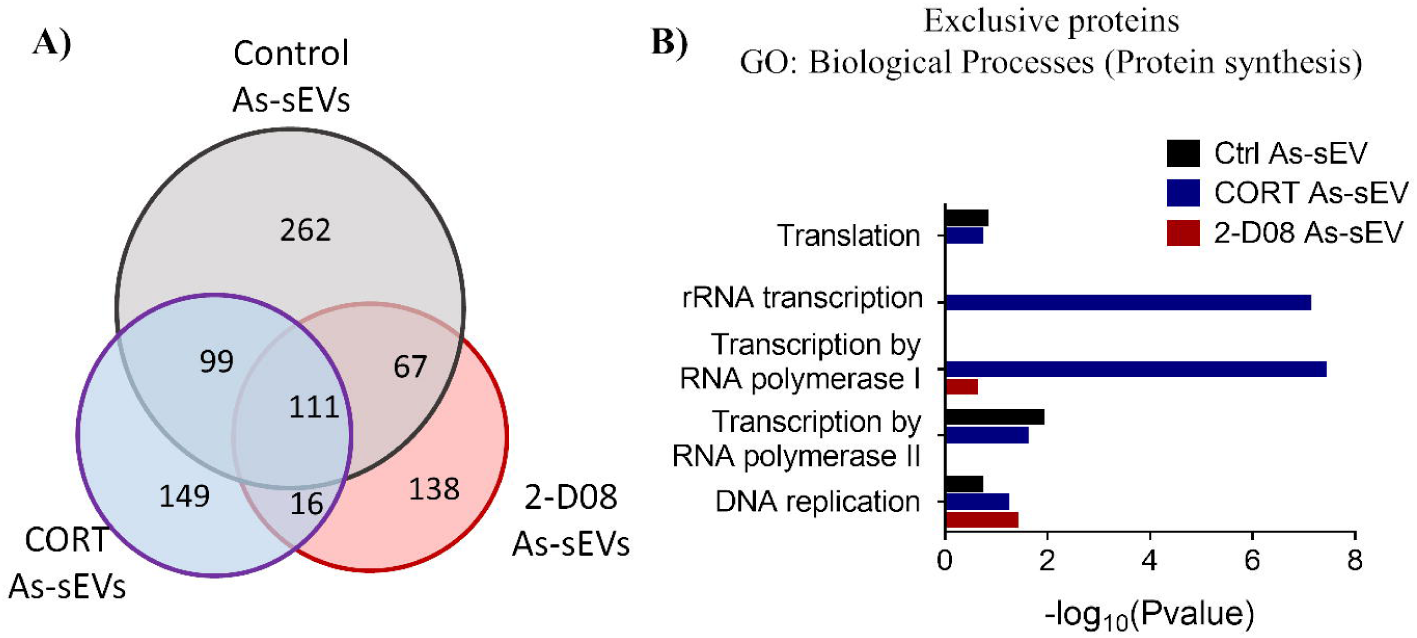
Proteomics of small Extracellular Vesicles (sEVs) from astrocyte treated with corticosterone reveals proteins related with protein synthesis. A) Venn analysis of sEV proteins from astrocytes treated with DMSO (control), corticosterone (CORT) and SUMO inhibitor (2-D08), identified by mass spectrometry. B) Comparison between Biological process present in exclusive proteins in each sample analyzed by the DAVID database. Data obtained from n=4-6 sEVs from different murine preparation.

Due to the large representation of protein synthesis regulatory proteins in sEVs after corticosterone treatment, we decided to study whether astrocyte-derived sEVs could modulate protein synthesis in target cells, i.e. in neurons. It was previously shown that astrocyte sEVs can be taken up by neurons (Luarte et al., 2020). Moreover, the relationship between SUMOylation and gene expression or transcription regulation has been described extensively (Liu & Shuai, 2008; Müller, Ledl, & Schmidt, 2004; Rosonina, Akhter, Dou, Babu, & Sri Theivakadadcham, 2017), as well as its relationship with protein translation (Hendriks & Vertegaal, 2016; Nie, Xie, Loo, & Courey, 2009; X. Xu, Vatsyayan, Gao, Bakkenist, & Hu, 2010).

Mature cortical neurons (21DIV) were incubated with sEVs derived from astrocytes treated with DMSO as a control, 2-D08 or CORT. To visualize the neurites of neurons in culture, immunofluorescence using a MAP2 antibody was performed and Sholl analysis was made to quantify dendritic arborization parameters (Figure 7A). We did not observe significant differences in the total dendrite length (Figure 7B). When analyzing the total number of intersections of dendritic branches (i.e. reflecting the dendritic arbor complexity), we found that neurons incubated with sEVs of control astrocytes presented a significant decrease compared to neurons that were not incubated with sEVs. This negative regulation is compatible with our previous results using 3 to 6 DIV neurons and suggest that astrocyte sEVs are implicated in dendritic pruning and/or in decreased growth (Figure 7C). Interestingly, an increase in the synthesis of new proteins in the soma of neurons incubated with sEVs derived from CORT-treated astrocytes was observed (Figure 7D and 7E). We observed no changes in the FUNCAT signal in neurons incubated with control astrocyte sEVs or with sEVs from astrocytes treated with the SUMOylation inhibitor (2-D08), compared with the signal in untreated neurons. This confirms that sEVs derived from corticosterone-treated astrocytes contain an increased number or level of proteins involved in the stimulation of protein synthesis.

**Figure 7.**
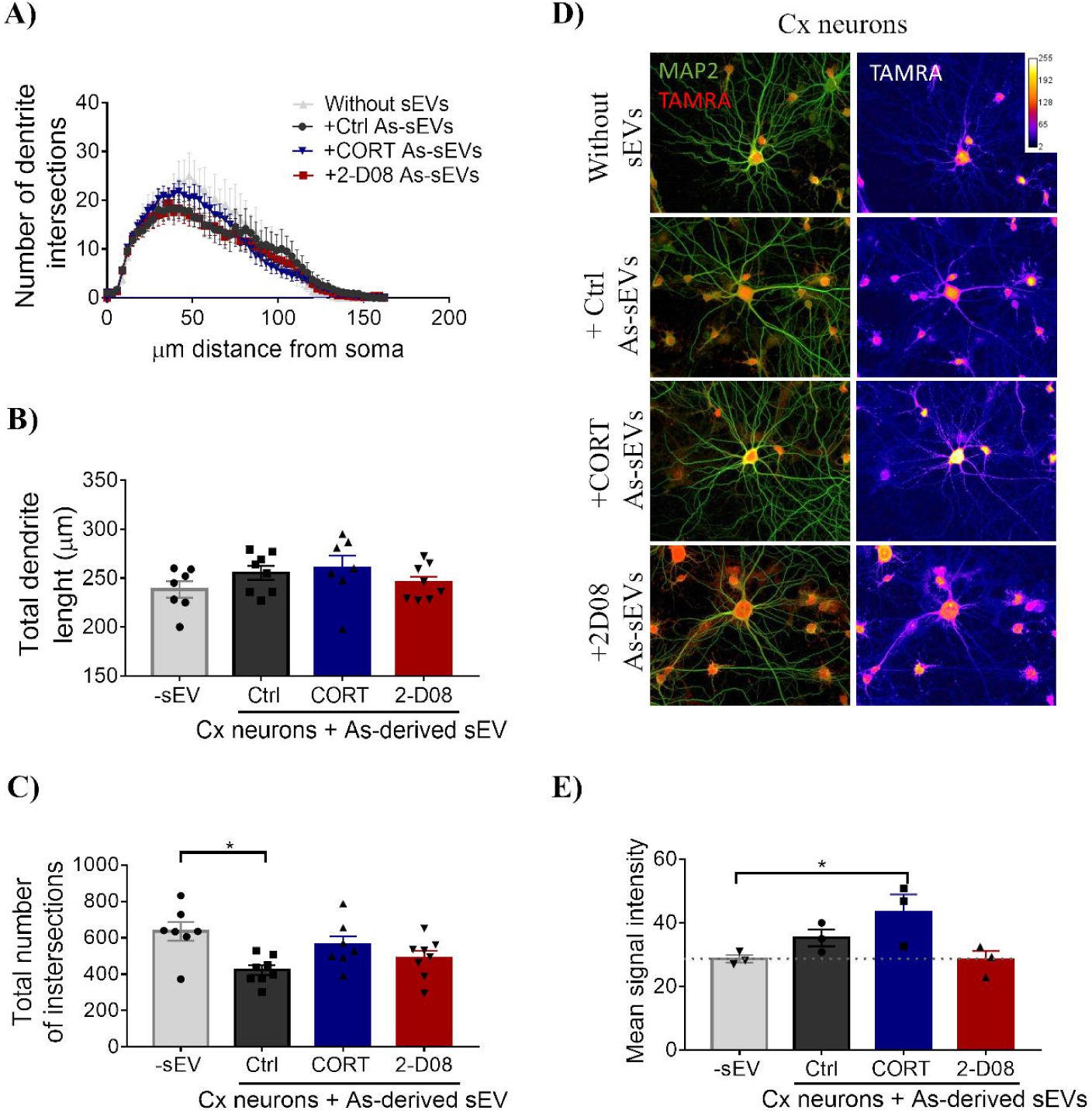
Neurons with small Extracellular Vesicles (sEVs) from astrocytes treated with corticosterone increase newly synthetized proteins. Cortical neurons untreated (-sEV) or incubated with 1000 astrocyte-derived sEVs per neuron (two times in 24 hour intervals). Astrocytes were previously treated with DMSO (Ctrl As-sEVs), corticosterone (CORT sEVs) and 2-D08 (2-D08 sEVs). 24 hours after the last incubation with sEVs, neurons were incubated with AHA or methionine added to the culture medium for 3 hours. The FUNCAT assay and MAP2 staining was performed. A) Sholl Analysis of neurons incubated with astrocyte-derived sEVs. B) Quantification of the total dendrite length per neuron. C) Quantification of total number of intersections per neuron. D) Neurons were stained with MAP2 antibody and newly synthetized proteins were detected by the AHA-TAMRA tag. The intensity of the FUNCAT signal is inserted as a fire lookup table in the right panel. E) Quantification of the intensity of the FUNCAT signal, TAMRA tagged-newly synthetized proteins (AHA). All data are expressed as mean ± SEM and for quantification one-way ANOVA was performed, followed by a post-hoc Tukey test. Statistical significance is represented with asterisks (n = 3-7, sEVs and cell culture from different murine preparation, 20 neurons was quantified by preparations *p<0.05).

## DISCUSSION

In recent years, sEVs have acquired an important place in the sights of many researchers in the area of neuroscience, due to the fact that they are secreted by most cells, can modulate target cell function (such as neuronal function), are present in the cerebrospinal fluid, and cross the blood brain barrier (Gómez-Molina et al., 2019; Liu et al., 2019). The presence of circulating sEVs in body fluids has opened the possibility that their molecular content could be used as a non-invasive diagnostic strategy to obtain biomarkers of physiological and pathological situations (Lin et al., 2015; Saeedi, Israel, Nagy, & Turecki, 2019; Zhang et al., 2019). Given the outstanding characteristics of these vesicles in the biomedical field, it was important for us to better understand the enigmatic mechanism of cargo biomolecule determination in them. Literature background led us to propose that SUMOylation corresponds to an important component that defines the proteins of sEVs.

### SUMOylation and its effect on astrocytes

In astrocytes, the overexpression of SUMO-1 and SUMO-2 did not substantially increase the levels of SUMOylation. This may be due to the low levels of transfection that we obtained in astrocytes, of about 15%. This is consistent with publications that describe the low transfection efficiency of astrocytes (Alabdullah et al., 2019), but also, that SUMOylation is a transient event that does not increase steady state levels (Henley, Craig, & Wilkinson, 2014; Klug et al., 2013).

In order to complement our observations regarding the role of SUMO overexpression, we used the SUMOylation inhibitor 2-D08. Other SUMOylation inhibitors such as Ginkgolic acid and Anacardic acid had been previously described (Fukuda et al., 2009), while more recently, the 2-D08 inhibitor was described (Kim et al., 2014). Several groups have already used this inhibitor to decrease SUMOylation: for example, in U937 cells, 2-D08 restored the anti-proliferative activity of retinoids (Baik et al., 2018; Lin et al., 2020; Lorente et al., 2019; Zhou et al., 2019). Accordingly, 2-D08 inhibited SUMOylation by SUMO-1 or SUMO-2, although both proteins regulate different cellular pathways. Specific SUMO inhibitors act at the level of non-covalent SIM interactions, thus providing an indirect inhibition of SUMO-regulated proteins (Hughes et al., 2017).

Previously, we had found that in serum sEVs of rats, a possibly SUMOylated form of ALDOC increased under stress conditions (Gómez-Molina et al., 2019; Ramírez, 2017). The stress response might be positively associated with this post-translational modification, to trigger the incorporation of a different set of proteins in sEVs. Thus, we decided to explore the effect of *in vitro* stress on SUMO levels in our experimental model using corticosterone. Despite observing an increase in SUMO-2ylation in astrocytes treated with corticosterone, we did not observe a significant increase in SUMO-1ylation. It would be interesting to explore under other *in vitro* stress conditions such as oxidative stress, heat stress, hypoxia, starving, or others whether this preference is a general stress effect. Moreover, these results favor again the idea that SUMO-1 and SUMO-2 protein modifications exert different biological roles or functions.

### SUMOylation and its effect on sEV cargo

The idea that SUMOylation participates in the process of protein loading in sEVs, is supported by the work showing that hnRNPA2B1 (Villarroya-Beltri et al., 2013) and α-synuclein (Kunadt et al., 2015) sorting depend on SUMOylation. Here, we determined that this mechanism broadly affects the loading of proteins into sEVs both in HeLa cells as well as in astrocytes.

Few studies focus on the total protein content in sEVs and/or the number of sEVs released under different SUMOylation conditions. Ageta described that the ubiquitin-like 3 (UBL3)/membrane-anchored Ub-fold protein (MUB) UBL3 is necessary for protein sorting in sEVs. In serum sEVs of UBL3 knock-out animals, a strong reduction of the total protein content in vesicles was found when gels were loaded with the same volume of a rat serum derived sEV suspension (Ageta et al., 2018). In a different study in cells transfected with siRNAs targeting TSG101 and Hrs, which are proteins of the ESCRT system, a decrease in the number and the total amount of proteins contained in sEVs was induced (Smith, Jackson, & Schorey, 2015). However, in this study the number of sEVs were not related to the parent cell number, which is an important issue to be considered, because the number of cells can vary according to the treatment.

Despite the low levels of SUMO expression in astrocytes, the modification by SUMO-2 caused an increase in protein content in the derived vesicles. One of the possible explanations for such an effect is the “SUMO paradox”, in general only a small fraction of the SUMO modified proteins is detected at a particular time point. However, while SUMO modification is involved in dynamic processes, many proteins will be modified at a given time. Thus, the enhancement or inhibition of SUMOylation strongly impacts the biological outcome, this also known as the history of SUMO modification (Hay, 2005) i.e. in our case, even low levels of SUMOylation is sufficient to influence protein loading of sEVs (Geiss-Friedlander & Melchior, 2007).

We show for the first time that inhibition of SUMOylation using 2-D08 in both astrocytes and HeLa cells could significantly increase the number of harvested sEVs. The release of other vesicle types, such as of synaptic vesicles, is regulated by SUMOylation (Craig, Anderson, Evans, Girach, & Henley, 2015; Tang, Craig, & Henley, 2015). Interestingly, in pancreatic ß cells and synaptic vesicles in neurons, the SUMOylation of STXBP5 (Tomosyn1A), a protein related with docking and fusion of vesicles is necessary for its binding with STX1A (Syntaxin-1A) and suppression of exocytosis. In such a way, inhibition of STXBP5 SUMOylation leads to an increase of exocytosis (Ferdaoussi et al., 2017). These proteins are also involved in the fusion of MVBs to the plasma membrane to release intraluminal vesicles (Geerts et al., 2017; Zhang et al., 2006). We thus speculate that SUMOylation of this protein could also be involved in the endo-exocytosis balance of MVB favoring the exocytosis when SUMOylation is inhibited.

### Identification of sEV proteins derived from cells that over-express SUMO

The SUMOylated proteins contained in sEVs and the biological processes in which these proteins are involved have been poorly described. We found proteins related to the synthesis of proteins in sEVs from astrocytes that overexpressed SUMO-1 or SUMO-2. Previously, the relationship of SUMO with chromosome stabilization, DNA replication, mRNA splicing and transcription and translation in cells had already been described in cells (Hendriks & Vertegaal, 2016; Nie et al., 2009; X. Xu et al., 2010). Possibly SUMOylated proteins in astrocyte sEVs were detected after pull downs with SIM domains. From them, EF-2, a protein related with protein synthesis was enriched in sEVs from S1 and S2 astrocytes, strongly suggesting that SUMOylation is a cellular process that defines EF-2 loading. However, the protein itself is detected at the expected molecular weight and thus, is unlikely to be SUMOylated, possibly it could be loaded into sEVs by interaction with another SUMOylated protein that acts as a “carrier”.

In turn, ALDOA was a common protein in sEVs from astrocytes related with cellular metabolism. We detected this protein with high molecular weights in sEVs, 55kDa and over 130kDa. Both molecular weights could coincide with different SUMOylation forms, the first is a mono-SUMOylation and the second a polySUMOylation (Pichler et al., 2017). The 130kDa form is mostly detected in sEV of astrocytes transfected with SUMO-1. It would be interesting to mutate the possible K residues that are SUMO acceptors to demonstrate that ALDOA loading depends on SUMOylation.

### Functional effect of astrocyte-derived sEVs on neurons

When neurons were incubated with control astrocyte sEVs, a decrease of dendritic arbor complexity was observed. This can be interpreted as increased pruning and thus enhanced network maturation during this time period or decreased growth (Luarte et al., 2020). Intriguingly, sEVs derived from corticosterone treated astrocytes did not affect dendritic length or complexity, but stimulated protein synthesis. The functional roles of the newly synthetized proteins need to be determined in the future. As the dendritic architecture depends on many different signaling pathways and molecular mechanisms (Arikkath, 2012; Skelton, Poquerusse, Salinaro, Li, & Luikart, 2020; J. Xu et al., 2019; Ziegler & Tavosanis, 2019). The contribution of specific molecules and proteins contained in sEVs on it deserves many future studies.

The methodology used by us to detect de novo protein synthesis in neurons allows visualization of newly synthesized proteins with high sensitivity (Dieck et al., 2012). The effect of sEVs on protein synthesis was abolished when astrocytes had been treated with SUMOylation inhibitor 2-D08, suggesting that SUMOylation is necessary to load proteins related with protein synthesis in sEVs. The relationship between SUMOylation and with chromosome stabilization, DNA replication, mRNA splicing and transcription has been widely described in cells (Hendriks & Vertegaal, 2016; Nie et al., 2009; X. Xu et al., 2010).

In summary when astrocytes are under stress, they increase the conjugation of proteins with SUMO-2 and allow the loading of proteins related to the synthesis of proteins within their sEVs. To confirm that SUMOylation is necessary for protein loading in sEVs, by inhibiting of SUMOylation with 2-D08 in 2 different cell types (HeLa and astrocytes) protein load per sEV was decreased, confirming the importance of SUMOylation in protein loading in sEVs. In turn, sEVs can cause changes in target cells such as neurons by changing dendritic arborization and increasing *de novo* protein synthesis, when the sEVs are from stressed astrocytes.

## Acknowledgments

We wholeheartedly thank Evelyn Dankert and Soledad Sandoval for her technical support and Dr. Holly Garringer and Dr. Grace Hallinan for critically reading the manuscript.

## Author Contribution

AF, AR and UW designed the experiments and wrote the manuscript. AF prepared all figures of the manuscript. The experimental work was done by: Figure 1, generated by AF; Figure 2, generated by AF; Figure 3, generated by AF and OS; TK supervised the mass spectrometry; AF did the bio-informatic analysis. Figure 4, generated by AF and MM; Figure 5, generated by AF. Figure 6 generated by AF; TK supervised the mass spectrometry; AF did the bio-informatic analysis. Figure 7 generated by AF; PL supervised the FUNCAT. Supplementary Figure 1 generated by AF. Supplementary Figure 2 generated by AF. Supplementary Figure 3 generated by AF. KC and TG prepared astrocytes primary cultures in all experiments.

## SUPPLEMENTARY FIGURE LEGENDS

**Supplementary figure 1. Transient transfection of SUMO plasmids in primary cultures of astrocytes has no effect on SUMO-1 or SUMO-2 expression**. A) Proteins conjugated with SUMO-1 or SUMO-2 were detected by Western blot in cells transfected with GFP, SUMO-1 (S1) or SUMO-2 (S2). The corresponding Coomassie staining is shown. B) Quantification of the proteins conjugated with SUMO-1 or C) SUMO-2 expressed as fold change with respect to the control (GFP). All data are expressed as mean ± SEM and for quantification one-way ANOVA was performed, followed by a post-hoc Tukey test. n=3, cell culture from different murine preparation.

**Supplementary figure 2. ALDOA and EF-2 are contained in astrocyte-derived small Extracellular Vesicles (sEVs)**. A) ALDOA, EF-2 and FLOT1 were detected by Western blot in sEVs from astrocytes transfected with GFP, SUMO-1 (S1) or SUMO-2 (S2). The same number of sEVs was loaded per lane. B) Quantification of ALDOA 55kDa double band in sEVs. C) Quantification of ALDOA 250kDa form. D) EEF-2 in astrocyte-derived sEVs. Fold change with respect to control (GFP). All data are expressed as mean ± SEM and for quantification one-way ANOVA was performed, followed by a post-hoc Tukey test. Statistical significance is represented with asterisks (n = 3, sEVs and cell culture from different murine preparation. *p<0.05).

**Supplementary figure 3. SUMOylation inhibition prevents proteins loading in small Extracellular Vesicles (sEVs) of HeLa cells**. Cells were treated 3 times with DMSO (control), 10 ng/ml IL-1β, 1μM corticosterone (CORT) or 30 μM 2-D08 (UBC9 inhibitor), in 24-hour intervals. sEVs were prepared 24 hours after the last treatment. A) Coomassie staining shows the sEV proteins contained in 10 million sEVs. B) Mean sEV size in nm. C) Number of vesicles released per million cells. D) Quantification of protein content (in ng) per sEV. All data are expressed as mean ± SEM and for quantification one-way ANOVA was performed, followed by a post-hoc Tukey test. Statistical significance is represented with asterisks (n = 9, sEVs and cell culture from different murine preparation, 3 technical repeats for each n. *p<0.05, ****p<0.001).

## SUPPLEMENTARY TABLE LEGENDS

**Supplementary Table 1**: List of proteins from astrocyte-derived sEVs. The columns contain the protein name, the Uniprot ID and in the following columns, the number of times that each protein was detected in sEVs samples from astrocytes transfected with GFP, SUMO-1 (S1) or SUMO-2 (S2). n=2 determinations for each case. Mass spectrometry was performed in Otto von Guericke University Magdeburg, Germany.

**Supplementary Table 2**: List of proteins pulled down with SIM beads from astrocyte-derived sEVs. The columns contain the protein name, the Uniprot ID and in the following columns, the number of times that each protein was detected in the starting material (input) and in the corresponding pull downs (PD) from astrocyte-derived sEVs. n=3 determinations for each case. Mass spectrometry was performed in Otto von Guericke University Magdeburg, Germany.

**Supplementary Table 3**: List of proteins from astrocyte-derived sEVs. The columns contain the protein name, the Uniprot ID and in the following columns, the number of times that each protein was detected in sEVs samples from astrocytes treated with DMSO (control), corticosterone (CORT) or 2-D08 (SUMOylation inhibitor). n=4-6 determinations for each case. Mass spectrometry was performed in Otto von Guericke University Magdeburg, Germany.

## REFERENCES

Ageta, H., Ageta-Ishihara, N., Hitachi, K., Karayel, O., Onouchi, T., Yamaguchi, H., … Tsuchida, K. (2018). UBL3 modification influences protein sorting to small extracellular vesicles. Nature Communications, 9(1). https://doi.org/10.1038/s41467-018-06197-y

Ageta, H., & Tsuchida, K. (2019, December 1). Post-translational modification and protein sorting to small extracellular vesicles including exosomes by ubiquitin and UBLs. Cellular and Molecular Life Sciences, vol. 76, pp. 4829–4848. https://doi.org/10.1007/s00018-019-03246-7

Alabdullah, A. A., Al-Abdulaziz, B., Alsalem, H., Magrashi, A., Pulicat, S. M., Almzroua, A. A., … Al-Mubarak, B. R. (2019). Estimating transfection efficiency in differentiated and undifferentiated neural cells. BMC Research Notes, 12(1), 225. https://doi.org/10.1186/s13104-019-4249-5

Arikkath, J. (2012, December 8). Molecular mechanisms of dendrite morphogenesis. Frontiers in Cellular Neuroscience. https://doi.org/10.3389/fncel.2012.00061

Baik, H., Boulanger, M., Hosseini, M., Kowalczyk, J., Zaghdoudi, S., Salem, T., … Bossis, G. (2018). Targeting the sumo pathway primes all-trans retinoic acid–induced differentiation of nonpromyelocytic acute myeloid leukemias. Cancer Research, 78(10), 2601–2613. https://doi.org/10.1158/0008-5472.CAN-17-3361

Colombo, M., Raposo, G., & Théry, C. (2014). Biogenesis, Secretion, and Intercellular Interactions of Exosomes and Other Extracellular Vesicles. Annual Review of Cell and Developmental Biology, 30(1), 255–289. https://doi.org/10.1146/annurev-cellbio-101512-122326

Craig, T. J., Anderson, D., Evans, A. J., Girach, F., & Henley, J. M. (2015). SUMOylation of Syntaxin1A regulates presynaptic endocytosis. Scientific Reports, 5, 17669. https://doi.org/10.1038/srep17669

Desterro, J. M. P., Rodriguez, M. S., Kemp, G. D., & Ronald T, H. (1999). Identification of the enzyme required for activation of the small ubiquitin-like protein SUMO-1. Journal of Biological Chemistry, 274(15), 10618–10624. https://doi.org/10.1074/jbc.274.15.10618

Desterro, J. M., Thomson, J., & Hay, R. T. (1997). Ubch9 conjugates SUMO but not ubiquitin. FEBS Letters, 417(3), 297–300. Retrieved from http://www.ncbi.nlm.nih.gov/pubmed/9409737

Dieck, S. T., Müller, A., Nehring, A., Hinz, F. I., Bartnik, I., Schuman, E. M., & Dieterich, D. C. (2012). Metabolic labeling with noncanonical amino acids and visualization by chemoselective fluorescent tagging. Current Protocols in Cell Biology, 1(SUPPL.56). https://doi.org/10.1002/0471143030.cb0711s56

Doyle, L., & Wang, M. (2019). Overview of Extracellular Vesicles, Their Origin, Composition, Purpose, and Methods for Exosome Isolation and Analysis. Cells, 8(7), 727. https://doi.org/10.3390/cells8070727

Ferdaoussi, M., Fu, J., Dai, X., Manning Fox, J. E., Suzuki, K., Smith, N., … MacDonald, P. E. (2017). SUMOylation and calcium control syntaxin-1A and secretagogin sequestration by tomosyn to regulate insulin exocytosis in human ß cells. Scientific Reports, 7(1). https://doi.org/10.1038/s41598-017-00344-z

Fukuda, I., Ito, A., Hirai, G., Nishimura, S., Kawasaki, H., Saitoh, H., … Yoshida, M. (2009). Ginkgolic Acid Inhibits Protein SUMOylation by Blocking Formation of the E1-SUMO Intermediate. Chemistry and Biology, 16(2), 133–140. https://doi.org/10.1016/j.chembiol.2009.01.009

Geerts, C. J., Mancini, R., Chen, N., Koopmans, F. T. W., Li, K. W., Smit, A. B., … Groffen, A. J. A. (2017). Tomosyn associates with secretory vesicles in neurons through its N- and C-terminal domains. PLOS ONE, 12(7), e0180912. https://doi.org/10.1371/journal.pone.0180912

Geiss-Friedlander, R., & Melchior, F. (2007, December). Concepts in sumoylation: A decade on. Nature Reviews Molecular Cell Biology, vol. 8, pp. 947–956. https://doi.org/10.1038/nrm2293

Gómez-Molina, C., Sandoval, M., Henzi, R., Ramírez, J. P., Varas-Godoy, M., Luarte, A., … Wyneken, U. (2019). Small Extracellular Vesicles in Rat Serum Contain Astrocyte-Derived Protein Biomarkers of Repetitive Stress. The International Journal of Neuropsychopharmacology, 22(3), 232–246. https://doi.org/10.1093/ijnp/pyy098

Hajmrle, C., Ferdaoussi, M., Plummer, G., Spigelman, A. F., Lai, K., Manning Fox, J. E., & MacDonald, P. E. (2014). SUMOylation protects against IL-1β-induced apoptosis in INS-1 832/13 cells and human islets. American Journal of Physiology. Endocrinology and Metabolism, 307(8), E664–73. https://doi.org/10.1152/ajpendo.00168.2014

Hay, R. T. (2005, April 1). SUMO: A history of modification. Molecular Cell, vol. 18, pp. 1–12. https://doi.org/10.1016/j.molcel.2005.03.012

Hecker, C. M., Rabiller, M., Haglund, K., Bayer, P., & Dikic, I. (2006). Specification of SUMO1- and SUMO2-interacting motifs. Journal of Biological Chemistry, 281(23), 16117–16127. https://doi.org/10.1074/jbc.M512757200

Hendriks, I. A., & Vertegaal, A. C. O. (2016). A comprehensive compilation of SUMO proteomics. Nature Reviews Molecular Cell Biology, 17(9), 581–595. https://doi.org/10.1038/nrm.2016.81

Henley, J. M., Craig, T. J., & Wilkinson, K. A. (2014, October 1). Neuronal SUMOylation: mechanisms, physiology, and roles in neuronal dysfunction. Physiological Reviews, vol. 94, pp. 1249–1285. https://doi.org/10.1152/physrev.00008.2014

Holopainen, J. M., Angelova, M. I., & Kinnunen, P. K. (2000). Vectorial budding of vesicles by asymmetrical enzymatic formation of ceramide in giant liposomes. Biophysical Journal, 78(2), 830–838. https://doi.org/10.1016/S0006-3495(00)76640-9

Hughes, D. J., Tiede, C., Penswick, N., Tang, A. A. S., Trinh, C. H., Mandal, U., … Whitehouse, A. (2017). Generation of specific inhibitors of SUMO-1- and SUMO-2/3-mediated protein-protein interactions using Affimer (Adhiron) technology. Science Signaling, 10(505). https://doi.org/10.1126/scisignal.aaj2005

Isola, A., & Chen, S. (2016). Exosomes: The Messengers of Health and Disease. Current Neuropharmacology, 15(1), 157–165. https://doi.org/10.2174/1570159x14666160825160421

Jeppesen, D. K., Fenix, A. M., Franklin, J. L., Higginbotham, J. N., Zhang, Q., Zimmerman, L. J., … Coffey, R. J. (2019). Reassessment of Exosome Composition. Cell, 177(2), 428–445.e18. https://doi.org/10.1016/j.cell.2019.02.029

Johnson, E. S., & Blobel, G. (1997). Ubc9p is the conjugating enzyme for the ubiquitin-like protein Smt3p. The Journal of Biological Chemistry, 272(43), 26799–26802. Retrieved from http://www.ncbi.nlm.nih.gov/pubmed/9341106

Kalluri, R., & LeBleu, V. S. (2020, February 7). The biology, function, and biomedical applications of exosomes. Science, Vol. 367. https://doi.org/10.1126/science.aau6977

Khan, F. A., Pandupuspitasari, N. S., Huang, C. J., Hao, X., & Zhang, S. (2016). SUMOylation: A link to future therapeutics. Current Issues in Molecular Biology, 18(1), 49–56. https://doi.org/v18/49 [pii]

Kim, Keyser S., & Schneekloth, J. (2014). Synthesis of 2′,3′,4′-trihydroxyflavone (2-D08), an inhibitor of protein sumoylation. Bioorganic and Medicinal Chemistry Letters, 24(4), 1094–1097. https://doi.org/10.1016/j.bmcl.2014.01.010

Klug, H., Xaver, M., Chaugule, V. K., Koidl, S., Mittler, G., Klein, F., & Pichler, A. (2013). Ubc9 Sumoylation Controls SUMO Chain Formation and Meiotic Synapsis in Saccharomyces cerevisiae. Molecular Cell, 50, 625–636. https://doi.org/10.1016/j.molcel.2013.03.027

Kowal, J., Arras, G., Colombo, M., Jouve, M., Morath, J. P., Primdal-Bengtson, B., … Théry, C. (2016). Proteomic comparison defines novel markers to characterize heterogeneous populations of extracellular vesicle subtypes. Proceedings of the National Academy of Sciences of the United States of America, 113(8), E968–77. https://doi.org/10.1073/pnas.1521230113

Kunadt, M., Eckermann, K., Stuendl, A., Gong, J., Russo, B., Strauss, K., … Schneider, A. (2015). Extracellular vesicle sorting of α-Synuclein is regulated by sumoylation. Acta Neuropathologica, 129(5), 695–713. https://doi.org/10.1007/s00401-015-1408-1

Liang, Y. C., Lee, C. C., Yao, Y. L., Lai, C. C., Schmitz, M. L., & Yang, W. M. (2016). SUMO5, a novel poly-SUMO isoform, regulates PML nuclear bodies. Scientific Reports, 6. https://doi.org/10.1038/srep26509

Lin, J., Li, J., Huang, B., Liu, J., Chen, X., Chen, X. M., … Wang, X. Z. (2015). Exosomes: Novel Biomarkers for Clinical Diagnosis. Scientific World Journal, 2015. https://doi.org/10.1155/2015/657086

Lin, Wang Y., Jiang, Y., Xu, M., Pang, Q., Sun, J., … Xu, J. (2020). Sumoylation enhances the activity of the TGF-β/SMAD and HIF-1 signaling pathways in keloids. Life Sciences, 255. https://doi.org/10.1016/j.lfs.2020.117859

Liu, B., & Shuai, K. (2008, June). Regulation of the sumoylation system in gene expression. Current Opinion in Cell Biology, Vol. 20, pp. 288–293. https://doi.org/10.1016/j.ceb.2008.03.014

Liu, Bai X., Zhang, A., Huang, J., Xu, S., & Zhang, J. Role of Exosomes in Central Nervous System Diseases., 12 Frontiers in Molecular Neuroscience § (2019).

Lorente, M., García-Casas, A., Salvador, N., Martínez-López, A., Gabicagogeascoa, E., Velasco, G., … Castillo-Lluva, S. (2019). Inhibiting SUMO1-mediated SUMOylation induces autophagy-mediated cancer cell death and reduces tumour cell invasion via RAC1. Journal of Cell Science, 132(20). https://doi.org/10.1242/jcs.234120

Luarte, A., Henzi, R., Fernández, A., Gaete, D., Cisternas, P., Pizarro, M., … Wyneken, U. (2020). Astrocyte-Derived Small Extracellular Vesicles Regulate Dendritic Complexity through miR-26a-5p Activity. Cells, 9(4). https://doi.org/10.3390/cells9040930

Mathieu, M., Martin-Jaular, L., Lavieu, G., & Théry, C. (2019, January 1). Specificities of secretion and uptake of exosomes and other extracellular vesicles for cell-to-cell communication. Nature Cell Biology, vol. 21, pp. 9–17. https://doi.org/10.1038/s41556-018-0250-9

Melchior, F. (2000). SUMO—Nonclassical Ubiquitin. Annual Review of Cell and Developmental Biology, 16(1), 591–626. https://doi.org/10.1146/annurev.cellbio.16.1.591

Miranda, K. J., Loeser, R. F., & Yammani, R. R. (2010). Sumoylation and nuclear translocation of S100A4 regulate IL-1β-mediated production of matrix metalloproteinase-13. Journal of Biological Chemistry, 285(41), 31517–31524. https://doi.org/10.1074/jbc.M110.125898

Müller, S., Ledl, A., & Schmidt, D. (2004, March 15). SUMO: A regulator of gene expression and genome integrity. Oncogene, vol. 23, pp. 1998–2008. https://doi.org/10.1038/sj.onc.1207415

Nie, M., Xie, Y., Loo, J. A., & Courey, A. J. (2009). Genetic and proteomic evidence for roles of Drosophila SUMO in cell cycle control, Ras signaling, and early pattern formation. PLoS ONE, 4(6). https://doi.org/10.1371/journal.pone.0005905

Pichler, A., Fatouros, C., Lee, H., & Eisenhardt, N. (2017). SUMO conjugation – a mechanistic view. Biomolecular Concepts, 8(1), 13–36. https://doi.org/10.1515/bmc-2016-0030

Ramírez, J. P. (2017). Vesículas extracelulares derivadas de astrocitos y su contenido de miRNAs son biomarcadores de estrés en la circulación periférica e interactúan con el epitelio intestinal. Thesis Magister. Universidad de los Andes.

Rosonina, E., Akhter, A., Dou, Y., Babu, J., & Sri Theivakadadcham, V. S. (2017, August 8). Regulation of transcription factors by sumoylation. Transcription, vol. 8, pp. 220–231. https://doi.org/10.1080/21541264.2017.1311829

Saeedi, S., Israel, S., Nagy, C., & Turecki, G. (2019, December 1). The emerging role of exosomes in mental disorders. Translational Psychiatry, Vol. 9. https://doi.org/10.1038/s41398-019-0459-9

Simons, M., & Raposo, G. (2009). Exosomes – vesicular carriers for intercellular communication. Current Opinion in Cell Biology, 21(4), 575–581. https://doi.org/10.1016/j.ceb.2009.03.007

Skelton, P. D., Poquerusse, J., Salinaro, J. R., Li, M., & Luikart, B. W. (2020). Activity-dependent dendritic elaboration requires Pten. Neurobiology of Disease, 134, 104703. https://doi.org/10.1016/j.nbd.2019.104703

Smith, Jackson L., & Schorey, J. S. (2015). Ubiquitination as a Mechanism To Transport Soluble Mycobacterial and Eukaryotic Proteins to Exosomes. The Journal of Immunology, 195(6), 2722–2730. https://doi.org/10.4049/jimmunol.1403186

Smith, P. K., Krohn, R. I., Hermanson, G. T., Mallia, A. K., Gartner, F. H., Provenzano, M. D., … Klenk, D. C. (1985). Measurement of protein using bicinchoninic acid. Analytical Biochemistry, 150(1), 76–85. https://doi.org/10.1016/0003-2697(85)90442-7

Song, J., Durrin, L. K., Wilkinson, T. A., Krontiris, T. G., & Chen, Y. (2004). Identification of a SUMO-binding motif that recognizes SUMO-modified proteins. Proceedings of the National Academy of Sciences of the United States of America, 101(40), 14373–14378. https://doi.org/10.1073/pnas.0403498101

Tang, L. T. H., Craig, T. J., & Henley, J. M. (2015). SUMOylation of synapsin Ia maintains synaptic vesicle availability and is reduced in an autism mutation. Nature Communications, 6. https://doi.org/10.1038/ncomms8728

Théry, C., Amigorena, S., Raposo, G., & Clayton, A. (2006). Isolation and Characterization of Exosomes from Cell Culture Supernatants and Biological Fluids. Current Protocols in Cell Biology, 30(1), 3.22.1–3.22.29. https://doi.org/10.1002/0471143030.cb0322s30

Van Niel, G., D’Angelo, G., & Raposo, G. (2018, April 1). Shedding light on the cell biology of extracellular vesicles. Nature Reviews Molecular Cell Biology, vol. 19, pp. 213–228. https://doi.org/10.1038/nrm.2017.125

Villarroya-Beltri, C., Gutiérrez-Vázquez, C., Sánchez-Cabo, F., Pérez-Hernández, D., Vázquez, J., Martin-Cofreces, N., … Sánchez-Madrid, F. (2013). Sumoylated hnRNPA2B1 controls the sorting of miRNAs into exosomes through binding to specific motifs. Nature Communications, 4, 2980. https://doi.org/10.1038/ncomms3980

Witwer, K. W., & Théry, C. (2019). Extracellular vesicles or exosomes? On primacy, precision, and popularity influencing a choice of nomenclature. Journal of Extracellular Vesicles, 8(1), 1648167. https://doi.org/10.1080/20013078.2019.1648167

Xu, J., Du, Y. L., Xu, J. W., Hu, X. G., Gu, L. F., Li, X. M., … Xu, J. (2019). Neuroligin 3 Regulates Dendritic Outgrowth by Modulating Akt/mTOR Signaling. Frontiers in Cellular Neuroscience, 13, 518. https://doi.org/10.3389/fncel.2019.00518

Xu, X., Vatsyayan, J., Gao, C., Bakkenist, C. J., & Hu, J. (2010). Sumoylation of eIF4E activates mRNA translation. EMBO Reports, 11(4), 299–304. https://doi.org/10.1038/embor.2010.18

Zhang, Liu Y., Liu, H., & Tang, W. H. (2019, February 15). Exosomes: Biogenesis, biologic function and clinical potential. Cell and Bioscience, Vol. 9. https://doi.org/10.1186/s13578-019-0282-2

Zhang, W., Lilja, L., Mandic, S. A., Gromada, J., Smidt, K., Janson, J., … Meister, B. (2006). Tomosyn Is Expressed in-Cells and Negatively Regulates Insulin Exocytosis. Diabetes, 55(3), 574–581. https://doi.org/10.2337/diabetes.55.03.06.db05-0015

Zhao, Q., Xie, Y., Zheng, Y., Jiang, S., Liu, W., Mu, W., … Ren, J. (2014). GPS-SUMO: A tool for the prediction of sumoylation sites and SUMO-interaction motifs. Nucleic Acids Research, 42(W1). https://doi.org/10.1093/nar/gku383

Zhou, P., Chen, X., Li, M., Tan, J., Zhang, Y., Yuan, W., … Wang, G. (2019). 2-D08 as a SUMOylation inhibitor induced ROS accumulation mediates apoptosis of acute myeloid leukemia cells possibly through the deSUMOylation of NOX2. Biochemical and Biophysical Research Communications, 513(4), 1063–1069. https://doi.org/10.1016/j.bbrc.2019.04.079

Ziegler, A. B., & Tavosanis, G. (2019, July 1). Glycerophospholipids – Emerging players in neuronal dendrite branching and outgrowth. Developmental Biology, vol. 451, pp. 25–34. https://doi.org/10.1016/j.ydbio.2018.12.009

